# A note on measuring natural selection on principal component scores

**DOI:** 10.1101/212258

**Authors:** Veronica K. Chong, Hannah F. Fung, John R. Stinchcombe

## Abstract

Measuring natural selection through the use of multiple regression has transformed our understanding of selection, although the methods used remain sensitive to the effects of multicollinearity due to highly correlated traits. While measuring selection on principal component scores is an apparent solution to this challenge, this approach has been heavily criticized due to difficulties in interpretation and relating PC axes back to the original traits. We describe and illustrate how to transform selection gradients for PC scores back into selection gradients for the original traits, addressing issues of multicollinearity and biological interpretation. In addition to reducing multicollinearity, we suggest that this method may have promise for measuring selection on high-dimensional data such as volatiles or gene expression traits. We demonstrate this approach with empirical data and examples from the literature, highlighting how selection estimates for PC scores can be interpreted while reducing the consequences of multicollinearity.

---

> *“Unfortunately, the biological meaning of principal components or discriminant functions is not always obvious.”—Endler (1986, p 192)*
>
> *“… it is difficult to translate from selection on principal components to selection on the original traits.”—Brodie et al. (1995)*

Understanding how natural selection acts on correlated traits remains a fundamental challenge in evolutionary biology (Lande and Arnold 1983). Most ecologically important traits that influence an organism’s fitness are correlated with other features of their morphology, life history, and behavior. The canonical approach to estimating selection on correlated traits is selection gradient analysis (Lande and Arnold 1983). By regressing estimates of relative fitness on multiple traits, the direct effects of individual traits can be distinguished from the effects of selection on correlated traits. Widespread application of the Lande-Arnold framework has revolutionized our understanding of natural selection, and has given empiricists a straightforward tool for evaluating the fitness consequences of phenotypic variation.

One challenge to the Lande-Arnold approach, however, has remained especially stubborn: regressing fitness estimates on correlated traits can become problematic when traits exhibit multicollinearity (Mitchell-Olds and Shaw 1987). If the correlation between traits is high enough, it becomes impossible to reliably estimate their separate contributions to relative fitness. Multicollinearity is frequently diagnosed with a combination of variance inflation factors (VIF) or condition indices produced by most regression packages, with a frequent rule of thumb that a VIF greater than 10 indicates serious multicollinearity (Belsley et al. 1980; Neter 1989; see also O’Brien 2007). Typical recommendations for multicollinearity include larger sample sizes, dropping or summing variables (Lande and Arnold 1983; Mitchell-Olds and Shaw 1987), principal component analysis (Lande and Arnold 1983; see below), or alternative approaches like partial least squares. While these methods have valuable statistical properties, their widespread adoption has been limited. In many instances estimating selection on highly correlated traits simultaneously is the object of study, and thus traits cannot be dropped. For partial least squares, interpretability of the coefficients or the factors identified can remain problematic, as does their lack of correspondence to selection gradients that can be used to predict evolutionary responses.

The merits of measuring selection on principal component scores, in particular, has been debated since the advent of selection gradient analysis. Lande and Arnold (1983) themselves measured selection on PC scores when analyzing selection on the Bumpus (1899) house sparrows dataset. Their use of principal components was for data reduction when analyzing 9 traits with 49 female birds. They noted that while directional selection was detected on some of the original traits, they failed to detect directional selection on PC1 scores. One of the major advantages of selection gradient analysis—measuring and comparing the magnitude of direct selection on correlated traits—can be lost, as one obtains an estimate of selection on a composite trait that may not be the same in sign or statistical significance as the original traits. PC scores are constructed based on their patterns of variation and covariation, regardless of their importance for fitness, and can be hard to interpret biologically, especially when they encompass many traits (see Mitchell-Olds and Shaw 1987 for an extended discussion). For these interpretive reasons, selection estimates on PC scores have been criticized (e.g., Endler 1986; Mitchell-Olds and Shaw 1987; Conner 2007; Hunt et al. 2007).

In this Comment and Opinion, we present and illustrate a simple approach that circumvents the interpretation problem of measuring selection on principal components. By projecting selection estimates for PCs back into the original trait space, their interpretability is preserved. We illustrate this with an empirical example in the mouse ear cress, *Arabidopsis thaliana,* as well as examples from the literature where previous authors have presented selection estimates for PC scores.

## Projecting selection gradients for PCs back into trait space

For most incarnations, traits will have separate units, and we assume that PCA has been performed on the correlation rather than the covariance matrix. A vector of selection gradients for PC scores (i.e., regression coefficients of relative fitness on PC scores) can be projected into the original trait space by:

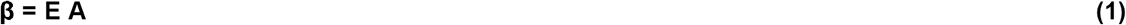

Where β is a vector of reconstituted selection gradients, **E** is a matrix with eigenvectors from the original PCA as columns, and **A** is the vector of regression coefficients obtained from regressing relative fitness on PC scores (Jolliffe 2002, pg 169). The coefficients in **A** describe the relationships between relative fitness and PC scores, while the elements of **E** describe the relationships between the PC scores and the original traits; combining them yields the relationship between traits and relative fitness. While an analogous equation for non-linear selection gradients would clearly be useful, we are unaware of any similar expression in the statistical literature.

Using all the eigenvectors of a PCA for **E** and regression with all PC scores as traits for **A** with (1) returns ordinary least squares regression coefficients (Jolliffe 2002 pg 169; Wilks 2005 pg 505), and does not eliminate multicollinearity. Using a subset of PCs can reduce multicollinearity, and estimate selection on the original traits. To estimate uncertainty in the elements of β, we estimate their variance using equation 6 of Lafi and Kaneene (1992), and from that standard errors. If one wishes to perform hypothesis testing on these reconstituted selection gradients, parameter estimates and standard errors could be used to estimate t-statistics and P-values. Prior to illustrating the use of (1) with our own data and literature examples, we note two important items that we return to below. First, using (1) necessarily involves a decision about how many PC axes to include, with attendant costs and benefits. Second, the elements of β estimated by (1) represent selection within the sub-space represented by the columns of **E**—e.g., if the columns of **E** represent 70% of the phenotypic variation, the elements of β represent selection on the traits in the sub-space describing 70% of the phenotypic variance.

## MATERIALS AND METHODS

### Size and phenology in *Arabidopsis thaliana*

The mouse ear cress, *Arabidopsis thaliana* (Brassicaceaae) is a selfing annual rosette plant. Our goal was to test the hypothesis that natural selection favoring early flowering, which is commonly observed in experiments even with low mortality (Austen et al. 2017), could be explained as selection on correlated traits, such as flowering duration and branch number. We reasoned that early flowering plants could potentially have longer flowering durations, and more opportunity to produce branches which could themselves support flowers and fruit. Stock (2015) crossed two parental lines collected from a single population in Doylestown, PA, (USA) that had alternative alleles at the flowering time genes *FRIGIDA (FRI)* and *FLOWERING LOCUS C (FLC;* Caicedo et al. 2004; Shindo et al. 2005) to produce an F1. She allowed this F1 to self-fertilize to produce F2 progeny, which have all allelic combinations at these two genes. Stock (2015) scored flowering time and allowed ∼500 F2 individuals to self-fertilize, producing F3 seed that we used for our experiment. We used five replicates from 50 F3 lines chosen to evenly span the F2 flowering time distribution.

We synchronized germination by stratifying seeds in 0.15 mg / 100 mL agar solution for six days at 4°C (Stock et al. 2015). We planted each seed in a standard conetainer with saturated Sunshine Mix #1 soil (Sun Gro Horticulture, Agawam, Massachusetts, USA). We arranged plants into conetainer racks in a randomized, blocked design, where the two shelves of a growth chamber represented experimental blocks, with plants of each line evenly divided between shelves. We maintained a light-temperature regime of 16 hour days (22°C) followed by nights of 20° C (Suter and Widmer 2013; Stock et al. 2015). We bottom-watered every two days by placing conetainers in trays of water for approximately three hours. After three months, we gradually reduced watering to simulate a terminal end of season drought, and all plants senesced after four months.

We scored bolting date, first and last flowering date throughout the experiment; at harvest, we measured branch number, rosette diameter, and counted the number of rosette leaves. We collected and counted fruits as a proxy for fitness (Stinchcombe et al. 2004). We measured flowering time as the number of days between planting and the first flowering date, and flowering duration as the number of days between first and last flowering dates. Analyses were performed using R v3.1.2 (R Foundation 2014) and SAS v9.4 (Cary, NC, USA).

### Trait correlations and selection

We calculated line means prior to analysis of correlations between traits and selection (Rausher 1992). We then used Pearson correlation coefficients as estimates of the line mean correlations among the traits. We first estimated selection differentials using simple univariate regressions regressing relative fruit number on individual traits, using line means and with traits standardized to a mean of zero and variance of one. We then estimated selection gradients for all traits, by regressing relative fruit number on them in a multiple regression. Strong multicollinearity between flowering time and duration lead to high variance inflation factors. To circumvent this, we used equation (1) above with four PCs, which explained 99% of the variation among line means.

### Literature examples

We searched for examples measuring selection on PCs to illustrate potential use of equation (1); we used published tables of PCAs and selection estimates for PC scores, rather than raw data. Our goal was to find illustrative examples rather than a comprehensive meta-analysis. We describe new findings obtained through the application of (1).

## RESULTS

### *Arabidopsis* experiment

The traits showed a range of correlations, ranging from rosette diameter and flower duration which were uncorrelated (r_g_ = 0.06, P = 0.647) to flowering time and duration, which were highly correlated (r_g_ = −0.95, P< 0.0001; Figure 1). Flowering time showed strong correlations with branch number (r_g_ = −0.74, P<0.001) and rosette leaf number (r_g_ =0.70, P <0.0001); flowering duration had essentially the opposite correlation pattern (r_g_ = 0.72 and r_g_ = −0.73, respectively, for branch number and rosette leaf number; Table 1). PCA showed that flowering time and flowering duration always had opposite signs in their loadings on PC1-PC4; rosette diameter, branch number, and rosette leaf number loaded heavily on PC2, PC3, and PC4.

**Table 1.**
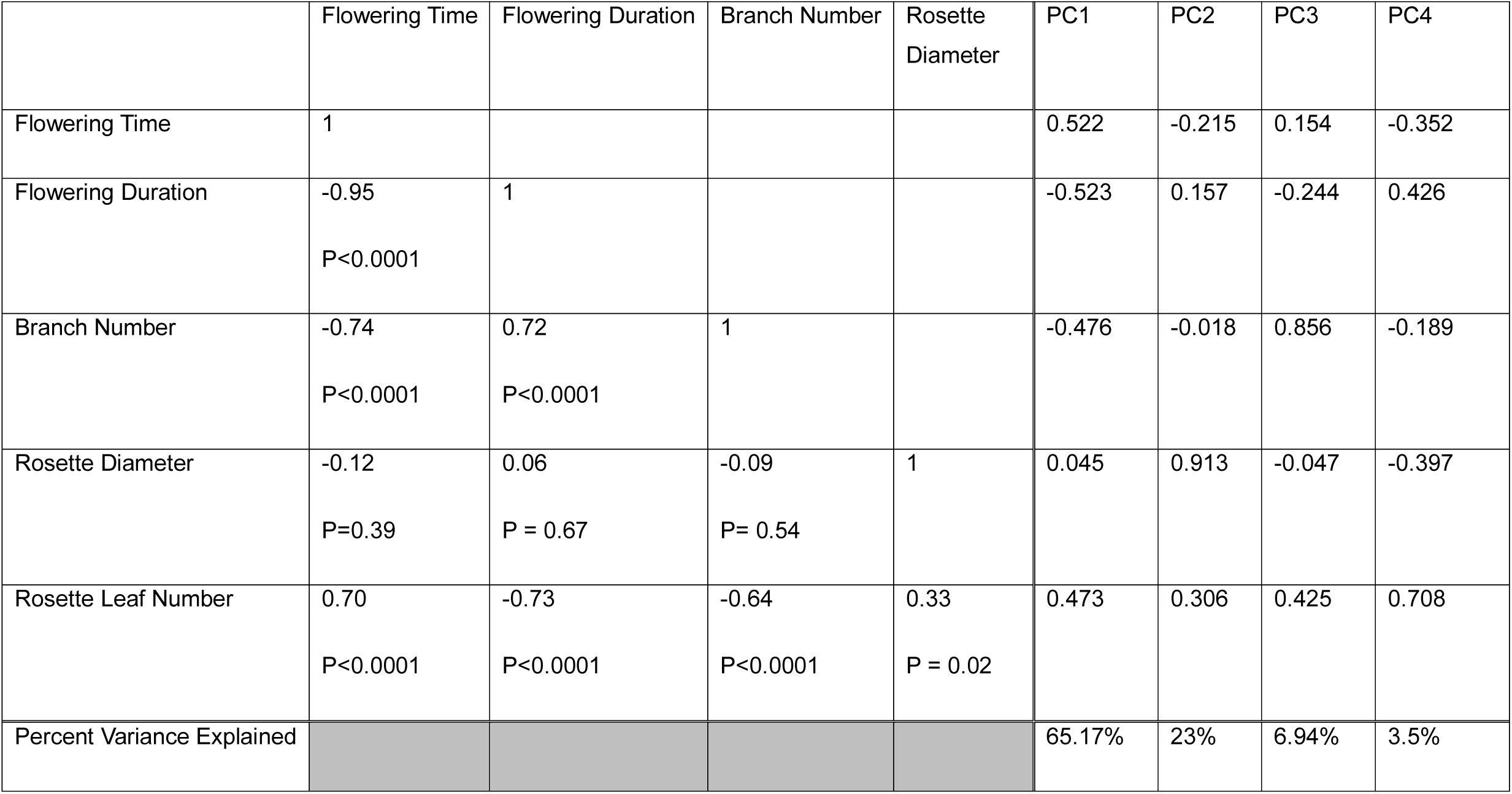
Line mean correlations for the *Arabidopsis* study, with eigenvector loadings for PC1-4.

|  | Flowering Time | Flowering Duration | Branch Number | Rosette Diameter | PC1 | PC2 | PC3 | PC4 |
| --- | --- | --- | --- | --- | --- | --- | --- | --- |
| Flowering Time | 1 |  |  |  | 0.522 | -0.215 | 0.154 | -0.352 |
| Flowering Duration | -0.95<br>P<0.0001 | 1 |  |  | -0.523 | 0.157 | -0.244 | 0.426 |
| Branch Number | -0.74<br>P<0.0001 | 0.72<br>P<0.0001 | 1 |  | -0.476 | -0.018 | 0.856 | -0.189 |
| Rosette Diameter | -0.12<br>P=0.39 | 0.06<br>P = 0.67 | -0.09<br>P= 0.54 | 1 | 0.045 | 0.913 | -0.047 | -0.397 |
| Rosette Leaf Number | 0.70<br>P<0.0001 | -0.73<br>P<0.0001 | -0.64<br>P<0.0001 | 0.33<br>P = 0.02 | 0.473 | 0.306 | 0.425 | 0.708 |
| Percent Variance Explained |  |  |  |  | 65.17% | 23% | 6.94% | 3.5% |

**Figure 1:**
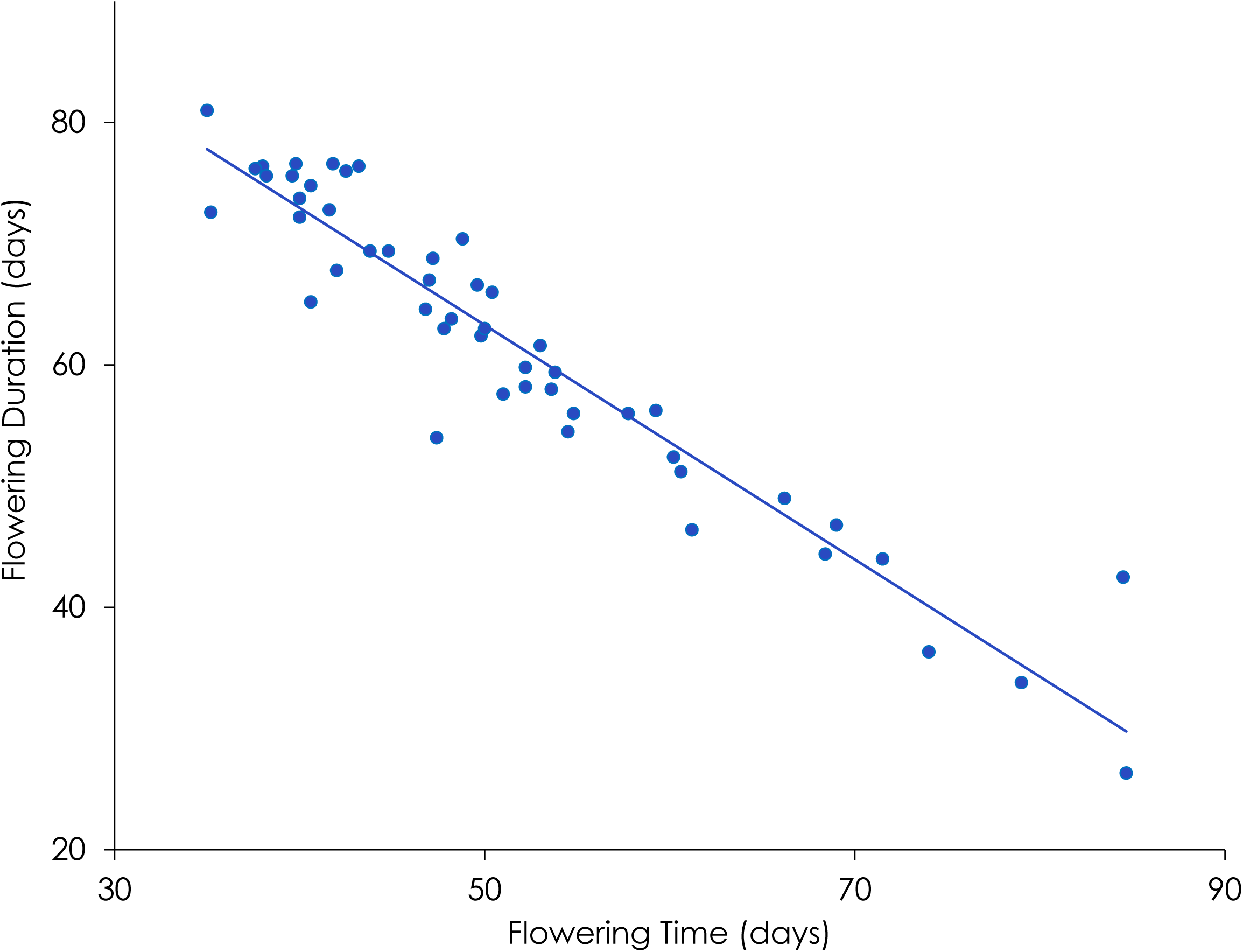
Line mean correlation between flowering time and flowering duration in the Arabidopsis chamber experiment. The inbred line mean correlation is −0.95.

### Selection on flowering time and flowering duration

Univariate estimates of selection suggest that early flowering, long flowering durations, more branches, and fewer rosette leaves are favored (Table 2a, Figure 2a-d). Interpreting these selection differentials is complicated by the strong correlations among traits: positive selection for branch number, for example, could be due to indirect selection for early flowering time (branch number is negatively correlated) or selection for longer flowering durations (branch number is positively correlated). Traditional multiple regression, while presented, is difficult to interpret: the high variance inflation factors for flowering time (12.54) and flowering duration (11.32) suggesting caution in interpreting these gradients. While both flowering time and flowering duration have large and significant selection differentials, neither are significant in the multiple regression, and both have standard errors approximately 3-4 fold larger than the standard error for the differentials, again because of multicollinearity.

**Table 2.** Selection estimates for the *Arabidopsis* experiment. Univariate estimates of selection are from regressions of relative fruit production on each trait alone. Selection gradients are estimated from a multiple regression of relative fitness on all traits, although the estimates for flowering time and duration are uncertain because of multicollinearity (indicated by variance inflation factors (VIF)). Principal component regression estimates are regression coefficients for PC1-PC4 projected back into the original trait space.

|  | Univariate Regressions |  |  | Multiple Regression |  |  |  | Principal Component Regression |
| --- | --- | --- | --- | --- | --- | --- | --- | --- |
| Trait | s ± se | t | P | $\beta$ ± se | t | P | VIF | $\beta \pm$ se |
| Flowering time | **-0.37**<br>±<br>**0.05** | **6.96** | **<0.0001** | -0.30<br>±<br>0.19 | 1.57 | 0.12 | 12.54 | -0.17 ± 0.05 |
| Flowering Duration | **0.35**<br>±<br>**0.06** | **6.29** | **<0.0001** | 0.05<br>±<br>0.18 | 0.27 | 0.78 | 11.32 | 0.17 ± 0.06 |
| Branch Number | **0.295**<br>±<br>**0.06** | **4.73** | **<0.0001** | 0.07<br>±<br>0.08 | 0.95 | 0.35 | 2.45 | 0.09 ± 0.08 |
| Rosette Diameter | 0.11<br>±<br>0.07 | 1.52 | 0.136 | 0.059<br>±<br>0.07 | 0.87 | 0.39 | 1.60 | 0.07 ± 0.07 |
| Rosette Leaf number | -<br>**0.218**<br>±<br>**0.07** | **3.18** | **0.0026** | 0.061<br>±<br>0.09 | 0.63 | 0.53 | 3.21 | -0.06 ± 0.10 |

**Figure 2:**
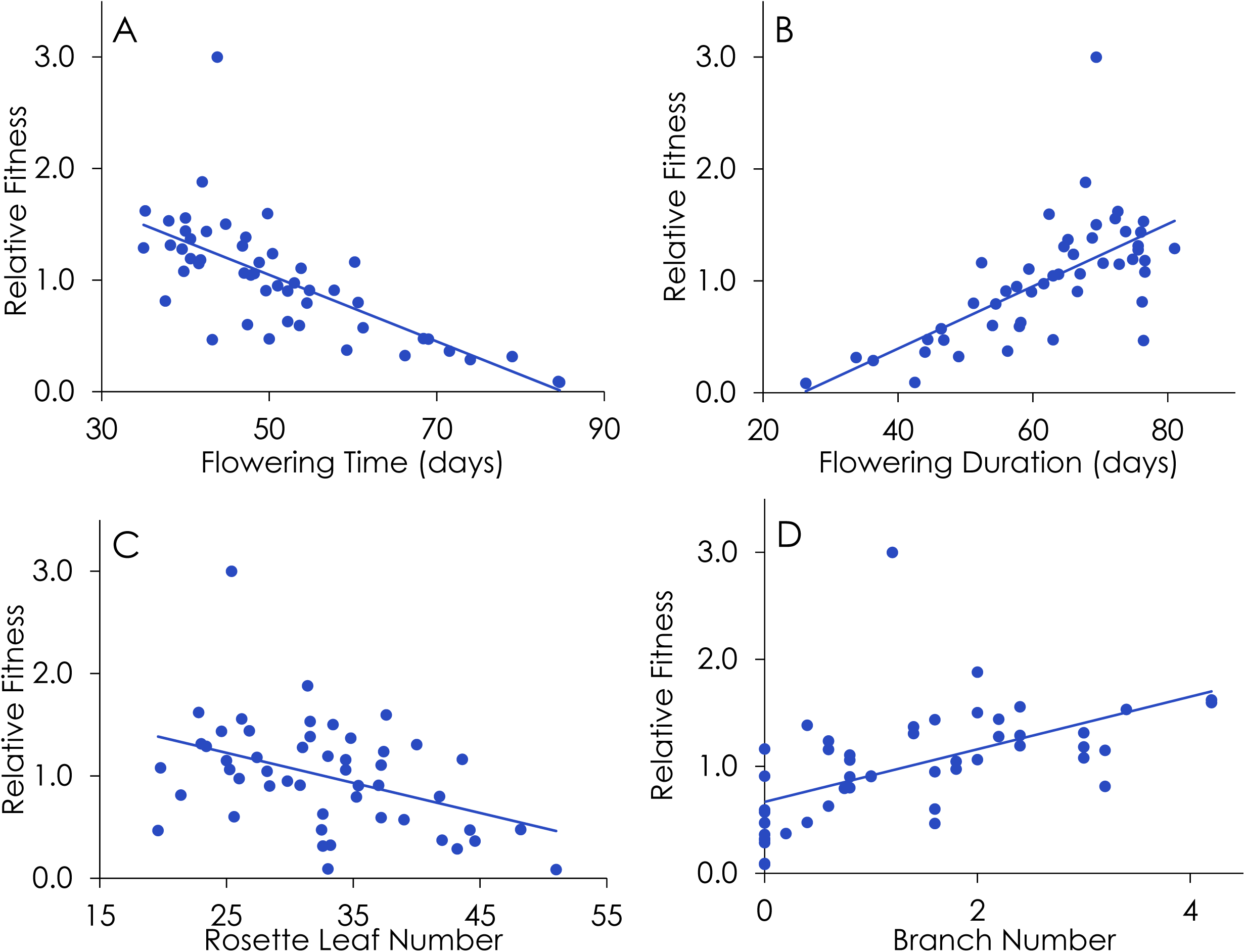
Selection differentials for: (A) flowering time and (B) flowering duration, (C) Rosette leaf number, and (D) Branch Number in the Arabidopsis experiment. Differentials are portrayed in the original trait units; plotted points are line means..

Estimating selection gradients via equation 1 using four PCs showed strong directional selection for earlier flowering (β ± s.e. = −0.17 ± 0.05), and for extended flowering durations (β *=* 0.17 ± 0.06). The sign of these selection gradients makes sense relative to their selection differentials and the sign and magnitude of the trait correlation; standard errors are approximately 1/3 to 1/4 of those observed in the multiple regression. These data suggest that selection is acting in approximately equal strength on flowering time and flowering duration, in opposite directions. The quantitative similarity of these selection gradients is likely due to the strong correlation between them, and reflects the two traits having approximately equal and opposite loadings on PC1-PC4 (Table 1). These data suggest that accounting for selection on correlated traits such as duration, branch number, size, and leaf number did not eliminate the observed selection favoring early flowering time, even if longer flowering durations are favored.

The estimated selection gradients for branch number, rosette diameter, and rosette leaf number are all approximately equal to their standard errors, suggesting that within the space of PC1-4, direct selection on these traits was weak. In addition, the reconstituted selection gradients for branch number, rosette diameter, and rosette leaf number (and their standard errors) are of roughly the same magnitude as in the original multiple regression, suggesting that calculations from (1) accurately characterized selection on them.

### Literature examples

#### Size and emergence in damselflies

Anholt (1991) measured selection on body size components in the damselfly *Enallagma boreale*. Several features of this study stand out. First, it was repeated across two years, and second, it included selection separately for males and females. Third, sample sizes were already large, and Anholt reported that he used PCs because of multicollinearity, as diagnosed by variance inflation factors. Anholt (1991) summarized the date of emergence, wing length, abdomen length, and mass at emergence with PCA, performing four separate PCAs, corresponding to each year-sex combination. He estimated selection differentials for PC1 and PC2 (which accounted for ˜85% of the variance, depending on sex-year combination), using both relative survival and relative mate acquisition as fitness estimates. We focus here on the survival data, as sample sizes were much larger (565 to 993 – vs-14 to 40). Because PC1 and PC2 are by definition uncorrelated, separate regressions of fitness on each PC to estimate selection differentials are equivalent to multiple regression of fitness on both PCs simultaneously. Consequently, we used the differentials for PC scores to calculate selection gradients.

Anholt interpreted PC1 as a size axis (due to the positive loadings of the size and mass characters), with the negative loadings of date of emergence on PC1 as reflecting the commonly observed trade-off between size and emergence date. In contrast, PC2 was interpreted as a date of emergence axis because date loaded most heavily, although wing length and abdomen showed moderate loadings in the same direction. As a consequence, PC1 represents a trade-off between date and size, while PC2 represents moderate positive associations between date and size. Because date of emergence figures in two PC axes in opposite directions vis a vis three other traits, and because the two PC axes were under selection in opposite directions in three of four cohorts, calculating selection gradients allows a straightforward assessment of the net pattern of selection on these traits without keeping track of the relative sign of multiple eigenvector elements and selection coefficients for PC scores across years and sexes. Our recalculations for the 1985 cohort suggest that selection in males favors early emergence dates (β = −0.35 ± 0.15) and large mass (β = 0.3321 ± 0.14), while in females the pattern is more variable on emergence date (β = −0.27 ± 0.19) and almost absent on mass (β = −0.065 ± 0.17). For the 1986 cohort, our calculations suggest that the dominant pattern in both males and females was for later emergence dates (β_males_ = 0.472 ± 0.18; β_females_ = 0.533 ± 0.26).

*Size traits in turtles:* Janzen (1993) studied selection on body size in hatchling snapping turtles (*Chelydra serpentine*). After releasing 112 hatchlings, he recaptured 66 individuals and estimated selection. Janzen summarized five size-related traits with PCA (initial egg mass, hatchling mass, carapace length, carapace width, and plastron length), using all of them as predictors of relative survival in a regression with other traits including locomotor performance, the size of the clutch that eggs came from, and water potential of the substrate eggs were incubated on (which affects size and hatching times). We used the first four PCs, which explained 99.1% of the phenotypic variance, and confined our attention to size traits used to construct the PCs. We note that subsequent work by Janzen (1998) suggests using logistic regression these cases, but we calculated selection gradients from multiple regression on PCs to illustrate the differences in interpretation.

When we projected selection estimates for the four PCs back into the original trait space, we found directional selection against hatchling mass (β = −0.360 ± 0.17), and in favor of greater carapace width (β = 0.370 ± 0.18) and plastron length (β = 0.309 ± 0.15). Two features of our reanalysis differ from the original results. First, Janzen reported positive directional selection on PC1, which he logically interpreted as an axis of size (because all elements of PC1 were positive for size-related traits). Our findings illustrate selection on the component traits of PC1 is not always in the same direction, even when all traits load on PC1 with similar sign and PC1 is under selection. The largest gradient we calculated was for decreased mass. The overall patterns of selection against mass, and in favor of carapace width and plastron length, come from selection on PC3 and PC4. PC3, which was under positive selection, had strong loadings of plastron length. PC4 had strong and nearly opposite loadings of hatchling mass and carapace width, which showed selection gradients of opposite sign when recalculated. While the overall pattern of selection on the individual traits make sense in light of selection on the PCs and the loadings of the overall traits, the converse is not true: without calculation, it is difficult if not impossible to infer net patterns of selection on individual traits from regression coefficients for PCs.

#### Tadpole morphology

Kraft et al. (2006) raised 560 tadpoles of the striped marsh frog (*Limnodynastes peroneii*) in either the presence or absence of dragonfly predators (*Anax* sp.), and measured the plastic response of morphological traits and their subsequent effects on survival. They measured nine morphological traits (total length, tail height, tail muscle height, body length, body height, tail area, tail muscle area, body width, and tail muscle width), and performed PCA on log transformed data. They found that PCs 4 and 5 (explaining 1.3% and 0.9% of the phenotypic variance, respectively) differed significantly between predator-induced and control tadpoles. They measured selection on all nine PCs in a multiple regression, finding significant negative selection on PC2, and positive selection on PC4 and PC5 for both the control and predator-induced individuals. Because sign is arbitrary within a PC axis, negative selection on PC2 is difficult to interpret: PC2 in their study had negative loadings of tail muscle width and positive loadings of body height. Individuals to the extreme of PC2 thus had either narrow tail muscles and large body heights, *or, equivalently*, wide tail muscles and small body heights. Which of these character combinations had lower fitness would make dramatic differences in the interpretation of selection.

When we projected selection coefficients for PC1-5 (representing 98.4% of the phenotypic variance), we found that selection was acting most strongly on body width in the control individuals (β = - 0.0395), while most strongly on tail muscle height in the induced individuals (β = 0.0426). (Standard errors for the regression coefficients for PCs were not presented, precluding estimation of standard errors of the reconstituted selection gradients). The case of tail muscle width and body height is also clarified: in both treatments, selection favors wider tail muscles and short body heights.

#### Flower volatiles and size

Schiestl et al. (2011) measured selection on 11 floral volatiles and inflorescence size in 70 individuals of the orchid *Gymnadenia odoratissima*. They performed several steps to facilitate selection analysis: first they reduced the number of floral volatiles analyzed from >40 to the 11 most abundant compounds in addition to inflorescence size. They summarized these variables with four principal components which were subjected to a varimax rotation, explaining 84% of the variation. The varimax rotation is meant to facilitate interpretation by creating a set of factors where each factor has predominant loadings from a reduced set of the original traits.

Projecting their selection coefficients for the factors back into the original trait space revealed appreciable selection on five measured traits: inflorescence size (β = 0.3208 ± 0.065), phenylacetaldehyde (β = 0.1580 ± 0.066), eugenol (β = 0.1290 ± 0.061), (z)-isoeugenol (β = −0.1733 ± 0.066), and phenylethyl (β = 0.1669 ± 0.061). Natural selection within the sub-space described by the first four PCs was approximately twice as strong on inflorescence size as on the volatile traits. Shiestl et al. (2011) detected negative selection on the third factor (most associated with α-pinene and (z)-isoeugenol after varimax rotation), yet α-pinene does not appear to be under selection when the coefficients are projected back into the original trait space.

#### Flower volatiles, display size, and phenology

Parachnowitsch et al. (2012) also measured selection on floral volatiles, in their case in the beardtongue *Penstemon digitalis*. They measured 30 traits in 88 individuals, and faced power challenges in estimating selection gradients for volatiles and other traits. Their solution was to only include traits in a multiple regression if they showed significant or marginally significant selection differentials. They noted several inherent challenges in measuring selection on floral volatiles: there are many compounds within a floral bouquet; simply measuring selection differentials cannot distinguish direct and indirect selection; if more than one trait loads on a PC, it is difficult to determine which trait is under selection if that factor is significant; and measuring selection on principal components or varimax-rotated factors inhibits comparisons to previous reports of selection on other floral traits.

We used the eight PCs presented by Parachnowitsch et al. (2012) which explained 72% of the variance. In addition to volatiles, they included morphological and life history traits, including flower size, corolla pigment, date of first flower, daily floral display size, number of days of flowering, and flower number. They observed very strong selection on PC3, which was associated with cis-6-nonenal, linalool, unknown-1, and daily display. By projecting selection estimates for eight PCs into the original trait space, we see that while selection was observed for the four traits associated with PC3 (β = 0.0569 ± 0.0167; β= 0.0468 ± 0.01286; β = 0.114 ± 0.0161; β = 0.1708 ± 0.01918, respectively), it was by far the strongest on daily display size. Interestingly, the one compound that was significant in the multiple regression performed by the authors, linalool, has an intermediate magnitude selection gradient when estimated in this manner.

## DISCUSSION

Our use of principal components regression combined with simple algebra offers the prospect of addressing the challenge of multicollinearity while solving the problems of interpreting selection on PC axes. Below we evaluate general trends that come from our reanalyses, compare the approach to related approaches in selection analysis, and close with a discussion of the limitations and judgements inherent to the approach we advocate.

### General lessons

We see three general lessons that have emerged from our approach. First, in numerous examples, it is much simpler to understand the reconstituted selection gradients on the original traits than on the PC axes, solving the interpretational challenges raised by Endler (1986), Mitchell-Olds and Shaw (1987), Brodie et al. (1995), Conner (2007), and Hunt et al. (2007). For example, for the *Arabidopsis* experiment, the net outcome of selection on flowering time and flowering duration being of approximately equal magnitude and opposite signs was difficult to see from the PC scores, and completely masked by the multiple regression. Likewise, in interpreting the Anholt and Kraft et al. studies, the original traits are straightforward to understand biologically, while selection on the PC axes can only be understood by considering multiple PC axes that have selection estimates of opposite signs, and are themselves made up of traits loading on those axes with opposite signs. While in some cases PCs have clear biological interpretations (e.g., the early flowering / small – vs-late flowering / large axis studied by Colautti et al. (2010] and Colautti and Barrett (2010]), for more complicated cases like the ones analyzed here projecting selection estimates on PCs back into the original trait space may be a valuable approach. It is important to recall, however, that these reconstituted selection gradients estimate how selection is acting on the portion of multivariate trait space spanned by the PCs that were originally included. Including more PCs captures more of the variation in the original traits measured, and is closer to the typical goal of selection gradient analysis.

Second, a natural interpretation of selection on PC axes is that they represent selection on the trait which loads most heavily on that axis. Several of our examples challenge this interpretation: for example, in the Schiestl et al. study, α-pinene loaded most heavily on an axis under selection, yet it did not appear to be under much selection when selection on PCs was projected back into the original trait space. Lande and Arnold (1983) noted that selection on PC scores can be diluted if traits under selection show strong phenotypic covariances with those unimportant for fitness. Our literature review revealed the converse problem: several cases in which seemingly straightforward selection on the PC axes was in fact a complex pattern of selection on the underlying traits. The reason for this is that traits load on all PC axes, rather than just one, and associations between these other PC axes and fitness contribute to the net pattern of selection on the traits.

Third, and perhaps most important, multicollinearity or high-dimensional data (e.g., floral volatiles, gene expression) do not need to be a stopping point for selection gradient analysis. For these types of studies, multicollinearity or high dimensionality will force investigators into doing *something*: either forgoing analysis, changing biological hypotheses, dropping traits, modifying their approach to selection gradient analysis, or facing complex interpretation challenges. The studies we reviewed were explicit about their approaches to these challenges. However, it is unclear how many studies in the literature encountered multicollinearity or high-dimensional data and dropped traits or changed their analysis approach, especially since the challenges and criticisms associated with measuring selection on PC axes may have convinced many authors to forgo PC regression. Measuring selection on PC axes and then projecting into the original phenotypic space, while not perfect, offers a route forward. We expect that PC approach may be useful for studies where investigators will typically have many more trait estimates (e.g., expression of many thousand genes, volatiles, metabolites, etc.) than individuals or estimates of relative fitness.

### Similarity to other approaches

Our approach is similar to other variants of selection gradient analysis, and it is worth considering the similarities and differences. At first glance, it appears similar to canonical rotation analysis (Simms 1990; Phillips and Arnold 1989; Blows 2007) because both involve PCA, although there are key differences. In that approach, the γ matrix of stabilizing, disruptive, and correlational selection gradients is subject to eigenanalysis, to identify the axes of trait variation under the strongest non-linear selection. In contrast, our approach is to apply PCA to the raw data, estimate linear selection on the synthetic axes, and then project back into the original trait space. Janzen and Stern (1998) presented a similar approach to ours, showing how to convert logistic regression coefficients into selection gradients that can be used to predict evolutionary change. Finally, our approach is somewhat similar to projection pursuit regression (Schluter and Nychka 1994). Projection pursuit regression can be applied to identify a single axis (a linear combination of traits), or multiple axes (multiple or combinations of traits) that explain the most variation in fitness (see also Morrissey and Sakrejda (2013) and Morrissey (2014)). It is important to note that projection pursuit regression rotates traits in a way that explains the most variation in relative fitness, while traditional PC axes are ordered by the variation in the traits themselves, but not fitness. Morrissey (2014) considered numerous types of regression methods for selection gradient estimation using both empirical data and simulations; he strongly supported projection pursuit regression. We see our approach as complementary, with one benefit being that it involves the typical machinery of least-squares regression and PCA familiar to many biologists, and can be applied retrospectively to summary data reported in papers (as we have here).

Our approach can also be interpreted in light of work on signal-to-noise ratios (Marroig et al. 2012). The similarity becomes clear when considering selection on PCs (recall that using equation 1 with all PC’s returns standard least squares estimates, so the following argument also applies to traits on the original scale). Trailing PCs represent a small amount of variation, and are estimated with greater error. Consequently, trailing PCs have a larger influence on inverted matrices, and because the role of matrix inversion at the heart of multiple regression, disproportionately contribute noise rather than signal to parameter estimates. In addition, the eigenvalues of leading PCs tend to be over-estimated, while those of trailing PCs are under-estimated (Kirkpatrick and Meyer 2008). Thus, even in the absence of any underlying correlation structure in the traits, noise and sampling error will have a dramatic effect on estimates of selection due to limited sample sizes (Marroig et al. 2012). Multicollinearity exacerbates this challenge by concentrating variation in a limited number of dimensions, making the estimation of trailing PCs, and hence selection on them, even more problematic for a given sample size. Marroig et al. (2012) illustrate alternative ways of dealing with signal to noise challenges for estimates of selection. They explore in detail an “extension” approach where unreliable eigenvalues representing sampling noise are replaced by the last reliable eigenvalue. In contrast, our approach eliminates the trailing eigenvalues and eigenvectors entirely, and measures selection in the phenotypic space described by the leading (and presumably most reliable) PCs. An important topic for future work might be to compare the extension method, our approach, as well as other forms of biased regression estimation (cf. Blair 2006; Morrissey 2014) for addressing the problems poised by multicollinearity and sampling variation.

### Prospects and caveats

Our use of PCA in combination with selection gradient analysis for our own data and examples from the literature entailed costs that are worth considering before wider application. First, the reconstituted regression coefficients are no longer unbiased estimators like least-squares estimates (Wilks 2005; Jolliffe 2002). By omitting PCs that explain relatively little variance, one reduces the variance in the estimated regression coefficients caused by multicollinearity (and their associated variance inflation factors), but at the expense of statistical bias (Jolliffe 2002). Jolliffe (2002, pages 170-171) shows how multicollinearity is driven by PCs with small eigenvalues, and accordingly, that regression coefficients for the predictor traits that load heavily on those axes will be estimated imprecisely in the original regression (see also Wilks 2005, page 505). Eliminating multicollinearity requires dropping a PC axis; it is important to note that information about all of the original traits still remains in the model, through the retained PC axes. In contrast, a trait omitted from a traditional regression could still have important effects on fitness, and be correlated with included predictors, but it is impossible to evaluate the effects of traits not included or reported in a selection analysis.

While omitting trailing PCs eliminates multicollinearity, Mitchell-Olds and Shaw (1987) criticized the assumption that trailing PCs (trait combinations with less variance) were not predictive of relative fitness and could be safely omitted. Likewise, Joliffe (1982) described several cases outside of evolutionary biology where PCs explaining relatively little variance were still predictive of the response variable. In the event that these PCs are associated with relative fitness, omitting them is not an advisable solution. In contrast, Blows and McGuigan (2015) and Sztepanacz and Blows (2017) show how the distribution of even *leading* eigenvalues from phenotypic and genetic data can be strikingly difficult to distinguish from sampling variation, suggesting that trailing principal components might primarily represent sampling variation. In the statistical literature, numerous approaches have been developed for determining how many PCs to use in principal component regression; Jolliffe (2002, pages 173-177) and Hill et al. (1977) provide overviews. One practical way forward would be to evaluate whether trailing PCs, which are responsible for multicollinearity, are predictive of relative fitness. If they are not (as in the *Arabidopsis* example), omitting these PCs and then projecting selection estimates back into the original trait space seems to be a viable option. Fundamentally, some judgement of the investigator will be required to decide how many PCs to include without reintroducing multicollinearity while still capturing a sufficient portion of phenotypic variance to evaluate biological hypotheses about selection.

## Acknowledgements

Our work is supported by NSERC Canada (JRS, VKC) and the Department of Ecology and Evolutionary Biology. We thank Ben Gilbert, Don Jackson, Dave Punzalan, Jessica Forrest, and Corlett Wood for discussion; comments by Anne Charmantier, Gabriel Marroig and two anonymous reviewers improved the manuscript.

## Data Accessibility

Empirical data and literature data are available on Dryad: <u>LINK.</u>

